# Exploratory 16S rRNA Metagenomic Analysis of Soil Microbial Communities in Agroecosystems of North-Central Argentina

**DOI:** 10.64898/2026.03.31.715494

**Authors:** Leticia Ana Guzmán, Cecilia Peralta, Antonella Marozzi, Eleodoro E. Del Valle, Leonardo Castoldi, Leopoldo Palma

**Author notes:** Corresponding author: Leopoldo Palma. Both authors contributed equally to this work (Leticia Ana Guzmán, Cecilia Peralta).

## Abstract

Agriculture has modified the soil structure due to the influence of external factors and processes that affect microbial biodiversity. Metagenomics is a fundamental tool for the study of soil microbial diversity because it provides information about the ecosystem diversity, including both the microorganisms that cannot be isolated in culture media and those that are no longer viable in the analyzed sample. In this work, six soil samples obtained from agroecosystems of central and northern Argentina were subjected to a preliminary 16S metagenomic analysis. Copiotrophic bacteria (Proteobacteria and Actinobacteriota) were dominant and one of the samples had a dominance of an oligotrophic Phylum (Acidobacteria). Our findings support previous evidence from traditionally managed agroecosystems and provide new insights into the diversity of soil microbiomes in Argentine regions outside the Pampas. Finally, we analyzed the most common genera with relevant species to agronomy, both beneficial and pathogenic, and their abundance and diversity in the sequenced samples.

## Introduction

The effects of anthropogenic activities are evident in landscapes and biological diversity, as well as at the intensification of erosion and climate change. These factors remark the need to modify production methods and their links with nature in order to promote a conscious and sustainable exploitation that prevents forcing the limits of soil capacity for providing its environmental services (Aizen et al., 2009; Guimarães, 1998). At the global scale, Argentina is the eighth largest country and has a low density of inhabitants, with population being concentrated in a few and quite focused spaces. Since the 20^th^ century, the country’s economy has been based mainly on exports of rural goods (Aizen et al., 2009), which are developed in more than 80% of its area and are related mainly to agricultural, livestock, and forestry practices (Bran et al., 2017).

The development of intensive agricultural practices can be analyzed at different scales, with the ecosystem-complexes scale being the minimum representation of the predominant landscape of an ecological region in Argentina (Aizen et al., 2009; Bran et al., 2017). This is an iterating approach, since modifications in them have been detrimental to environmental quality at all the other different scales. During the 1990-2006 period, a shift in the productive structure occurred towards soybean as the main Argentine crop, with a yield increase of approximately 45% (Aizen et al., 2009). The modifications that occurred in these territories are not only at the landscape scale but also at the level of the environmental services they provide, like the loss of soil quality, water absorption, and the alteration of biodiversity (Aizen et al., 2009).

Land use patterns have several effects on soil biodiversity, food webs, and ecosystem services (Bran et al., 2017). Agricultural activity has replaced the biological processes that regulate soil structure through external factors, such as the application of fertilizers or synthetic pesticides. In this sense, changes driven by different management practices alter soil properties, causing a direct effect on microbial communities, also known as soil microbiome. For example, modifications produced by fertilizers change the soil chemical properties that may have an influence over the presence and prevalence of some bacteria groups, preventing the growth of others (Dai et al., 2018). Furthermore, chemical fertilizers have different effects than those of organic fertilizers. Likewise, the overuse of nitrogen (N) causes the loss of organic carbon, with an impact on microbial biodiversity, that also increases nitrification and leaching of nitrate to nitrogen and reaching groundwater sources (Dai et al., 2018; Geisseler and Scow, 2014). Among the traditional practices, differences were also observed between fertilization with N alone and with N complemented with phosphorus (P) and potassium (K) ^(^Dai et al., 2018^)^. The modification of microbial communities due to the application of different agricultural practices has an impact on the agroecosystems, since microorganisms participate in soil formation through nutrient inputs (e.g. phosphorus and nitrogen), degradation of recalcitrant organic matter, and mineral weathering (Dubey et al., 2019). In this regard, a better understanding of the factors determining the structure of soil microbiomes may contribute to a better understanding of soil dynamics and change the approach of the most widely used traditional agricultural practices.

Microbial diversity should be considered an indicator of soil health and an important factor to ensure agriculture sustainability (Kraut-Cohen et al., 2020). Metagenomic analyses allow us to identify the microorganisms that cannot be isolated and grown in culture media; consequently, the information obtained using this method has become a fundamental tool to correlate factors that have an influence on soil microbiota in an evolving environment (Dubey et al., 2019). These analyses have been conducted in agricultural systems to assess variations in soil biodiversity relative to management type (Rascovan et al., 2013). In Argentina, some metagenomic studies were done in the Pampas agroecosystems (Fierer et al., 2013; Rascovan et al., 2013); other productive regions of the country, however, have not been subjected to these analyses.

In this work, we conducted an exploratory metagenomic analysis of the bacterial communities present in agricultural soil samples belonging to north-central region of Argentina, including the Chaco and Pampas ecosystem complexes.

## Material and methods

### Area of analysis

Soil samples were collected from different agroecosystems cultivated with soybean (*Glycine max* L.) and corn (*Zea mays* L.) in Argentina (Morello et al., 2012). The agroecosystems were in the three most populated ecoregions of the country under an extensive agricultural production that causes significant modifications of the landscape structures.

The Dry Chaco ecoregion is part of the Great South American Chaco and is characterized by a large flat terrain with a slight slope towards the east of Córdoba province. The climate conditions are widely varied, with the highest summer temperatures exceeding 47 Celsius degrees (°C), and the lowest winter temperatures dropping to below –6 °C. Cumulative volumetric rainfall ranges from 400 to 800 mm per year and predominant soil types include Mollisols, Entisols, and Alfisols. The landscapes of this ecoregion have also suffered different transformations after cotton cells were consolidated, technological changes of agriculturalization processes have been incorporated for soybean production and cattle raising in the areas where soybean cannot be cultivated. The soils are poorly enriched, with a slight tendency to be saline. In the Central Subhumid Chaco Complex, the physiognomy is dominated by halophytic savannahs and Chaco forest, which were fragmented by land use conversion to extensive agriculture (Morello et al., 2012) (Figure 1).

**Figure 1.**
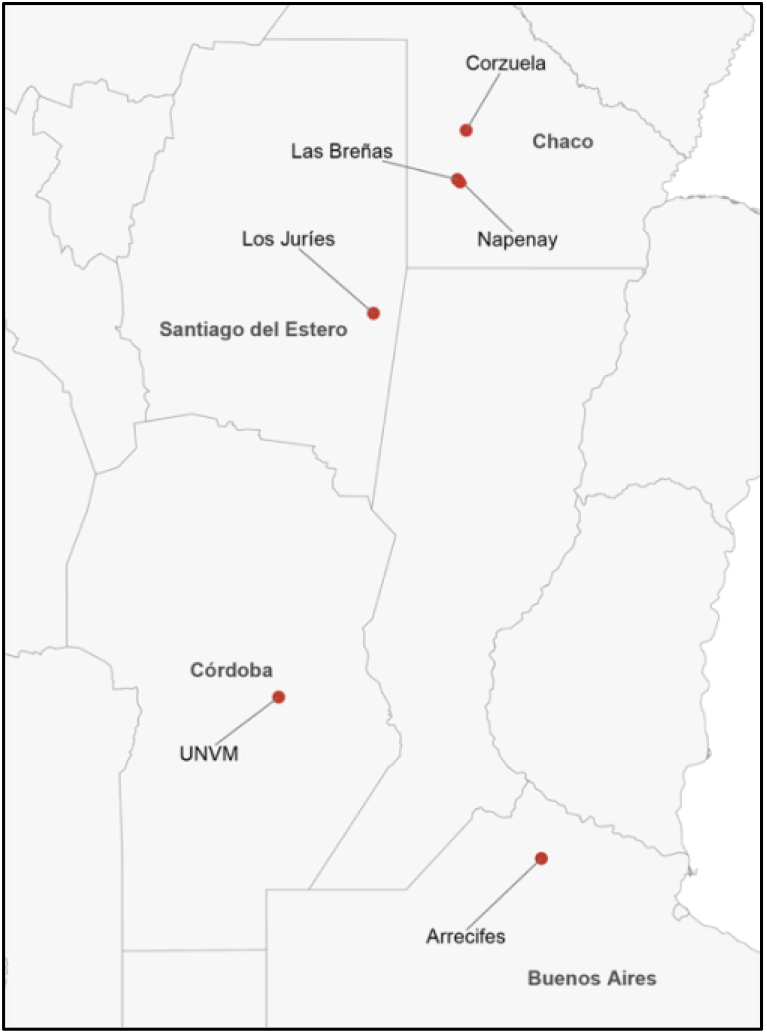
Geographic location of soil sampling sites in Argentina. Samples were collected from agroecosystems in the Chaco region (Corzuela, sample 69; Las Breñas, sample 70; Napenay, sample 71), Santiago del Estero (Los Juríes, sample 82), Córdoba (UNVM), and Buenos Aires (Arrecifes, sample 83). Points indicate sampling locations and province names are shown for geographic reference.

The Espinal ecoregion is represented by a flat to gently undulating terrain, including a lesser extent low mountain range and loessial to sandy soils, Mollisols, Vertisols, and Alfisols that stand out in the Northern Complexes, with a slightly more humid and warmer climate. Cumulative volumetric rainfall ranges between 500 and 1000 mm per year, decreasing from north to south. This ecoregion is characterized by the presence of low xerophytic forests, including alternating grasslands and savannas. The loss of the natural physiognomy in this ecoregion is one of the most important of the country and has occurred due to processes of anthropization associated with crop production and urbanization. This region includes the Pampa Pedemontana Complex, which is present almost entirely in Córdoba province (Morello et al., 2012).

The Pampas ecoregion is located the east-central region of Argentina and is an extensive plain with soft undulations in some areas and a slope from West to East towards the Atlantic Ocean. The predominant climate is subtropical, with a humidity gradient that decreases from East to West and Southwest; cumulative volumetric rainfall ranges from 1200 mm to 700 mm annually. Average annual temperature ranges from 20 to 14 °C, decreasing latitudinally. The predominant soils are Mollisols, mostly Argiudols, Haplustols, Hapludols, and Natracuols (Morello et al., 2012). The dominant vegetation type is the grass steppe, and the species composition varies according to the local climatic and soil characteristics. This ecoregion has the longest history of agricultural, and livestock use in the country and is divided into two sub-regions; our study was conducted in the Undulating Pampas complex, which is in the north and is predominantly a grassland. However, most of the area has been converted into farmlands, and the pastures were replaced with crops, except in some relictual spaces, where agricultural activity cannot be developed (Morello et al., 2012).

### Sample collection and analysis

Soil samples were collected using a tubular soil sampler from the top 20 cm of soil, as a composite of 10 random subsamples (∼100 g each). A total of six samples were obtained from agroecosystems under conventional agricultural management, predominantly Mollisols. Four samples were collected from the Central Subhumid Chaco Complex: Corzuela (sample 69), Las Breñas (sample 70), Napenay (sample 71), and Los Juríes (sample 82). The remaining two samples were collected from the Pampa Pedemontana (National University of Villa María, UNVM) and the Pampa Ondulada Complex (Arrecifes, sample 83). All samples were collected in February 2018 from soybean-cultivated soils, except for UNVM, which was under corn cultivation.

### DNA purification, 16S rRNA sequencing and analysis

Soil samples were homogenized in a high-speed mixer (High-speed Universal Disintegrator, Pro-Lab Diagnosis), and DNA extraction was performed using the PureLink Microbiome DNA Purification Kit (Invitrogen). DNA quality was assessed by electrophoresis on a 1% agarose gel using SYBR Safe DNA staining (Thermo Fisher Scientific), and concentration was measured using a PICODROP spectrophotometer (PICO 100).

Amplicon libraries targeting the V3–V4 regions of the 16S rRNA gene were constructed and sequenced using Illumina MiSeq technology (Macrogen). Raw reads were quality filtered, trimmed, and merged to obtain high-quality sequences. Reads shorter than 50 bp were removed.

Subsequent data processing and analysis were conducted in R. Taxonomic abundance tables were used to calculate diversity indices, including Shannon diversity and richness, using the vegan package (Oksanen et al., 2013). Relative abundance at different taxonomic levels (phylum and genus) was calculated from read counts.

Genus-level abundance data were aggregated and used to assess the distribution of agronomically relevant taxa. Genera were grouped based on their reported roles as plant-beneficial or plant-associated bacteria (Ali, 2019), while plant-pathogenic genera were identified according to the classification proposed by Mansfield et al. (2012).

Raw sequencing data were deposited in the NCBI Sequence Read Archive (SRA) (Leinonen et al., 2010) under BioProject accession number PRJNA722672.

## Results

### Microbial community structure at phylum level

The taxonomic composition of soil microbiomes revealed consistent dominance of a limited number of major bacterial phyla across all sampled sites. The most abundant phyla were Proteobacteria, Actinobacteriota and Acidobacteriota, generally exceeding 15% relative abundance in all samples (Fig. 2). Proteobacteria was the dominant phylum in Corzuela, Los Juríes, Arrecifes and UNVM, representing more than 40% of the total community in some cases. In contrast, Actinobacteriota predominated in Napenay, while Acidobacteriota showed its highest relative abundance in Las Breñas, suggesting differences in soil physicochemical conditions among sites. Other phyla, including Chloroflexi, Gemmatimonadota, Planctomycetota and Verrucomicrobiota, were consistently detected at lower abundance, indicating a complex and diverse microbial structure across all agroecosystems. The relative abundance of dominant bacterial phyla is shown in Figure 2.

**Figure 2.**
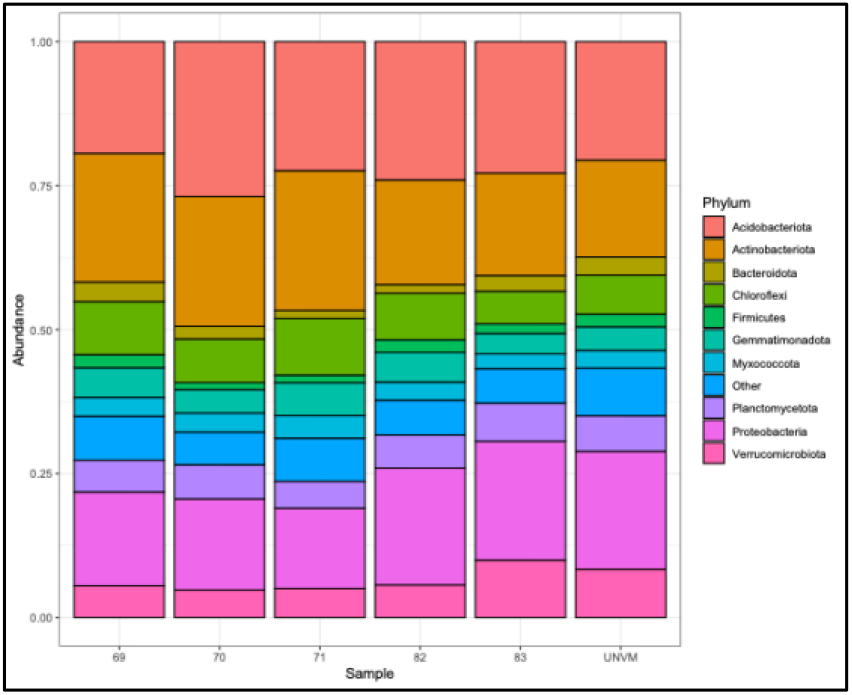
Relative abundance of dominant bacterial phyla across soil samples from different agroecosystems in north-central Argentina. Samples correspond to distinct geographic locations: 69 (Corzuela, Chaco), 70 (Las Breñas, Chaco), 71 (Napenay, Chaco), 82 (Los Juríes, Santiago del Estero), 83 (Arrecifes, Buenos Aires), and UNVM (Córdoba). Taxa are shown at the phylum level, and only the most abundant groups are displayed, with less abundant taxa grouped as “Other.”

### Alpha diversity and richness

All soil samples exhibited high bacterial diversity. Shannon index values ranged from 7.46 to 7.77, with the highest diversity observed in Arrecifes and UNVM, and the lowest in Los Juríes, suggesting potential local differences in community evenness (Figure 3A). Observed richness ranged from approximately 2610 to 3424 amplicon sequence variants (ASVs), indicating the presence of highly complex microbial communities in all sampled soils (Figure 3B).

**Figure 3.**
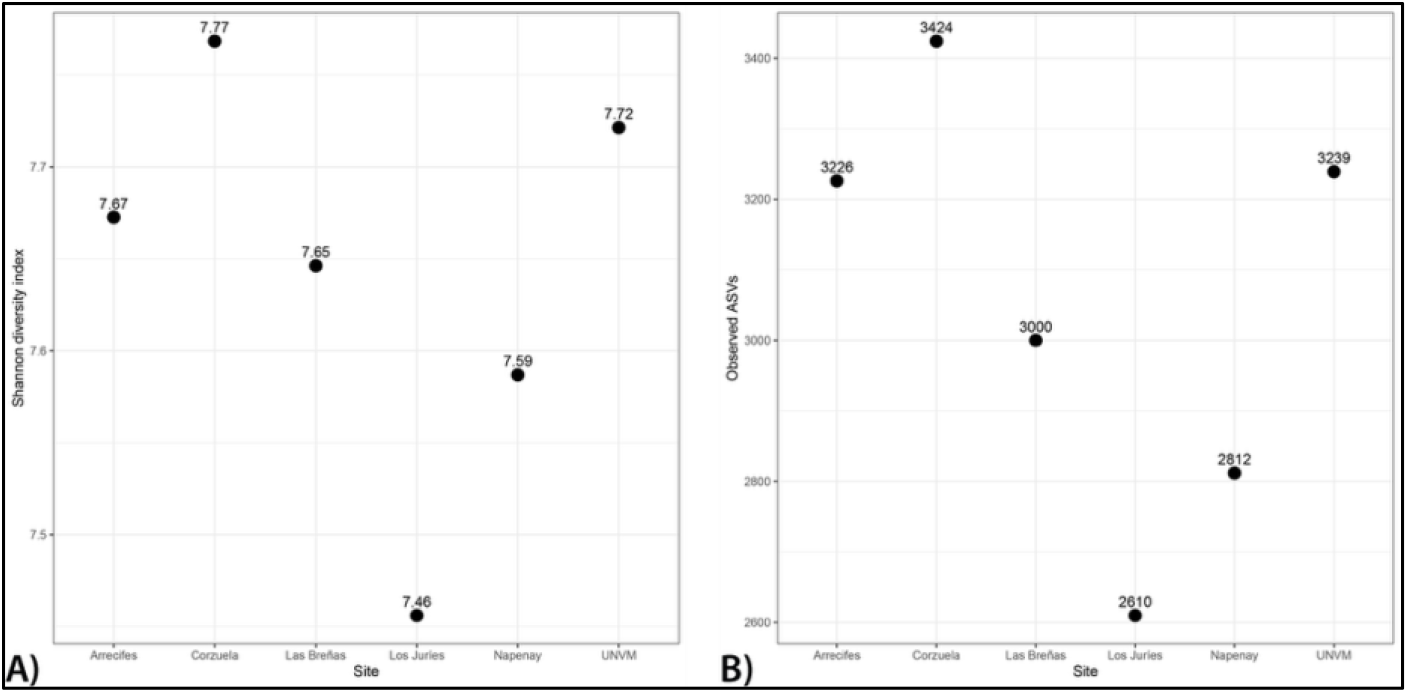
Alpha diversity of soil bacterial communities across sampled agroecosystems. (A) Shannon diversity index and (B) observed amplicon sequence variants (ASVs) for each sampling site.

Although variation in Shannon diversity and ASV richness was observed across sites, these differences were not statistically evaluated due to the limited number of samples.

### Beta diversity and community differentiation

Principal coordinate analysis (PCoA) based on Bray–Curtis dissimilarity revealed clear differentiation among sampling sites, with the first two axes explaining a substantial proportion of the variance, indicating clear differences in community structure across sites (Fig. 4).

**Figure 4.**
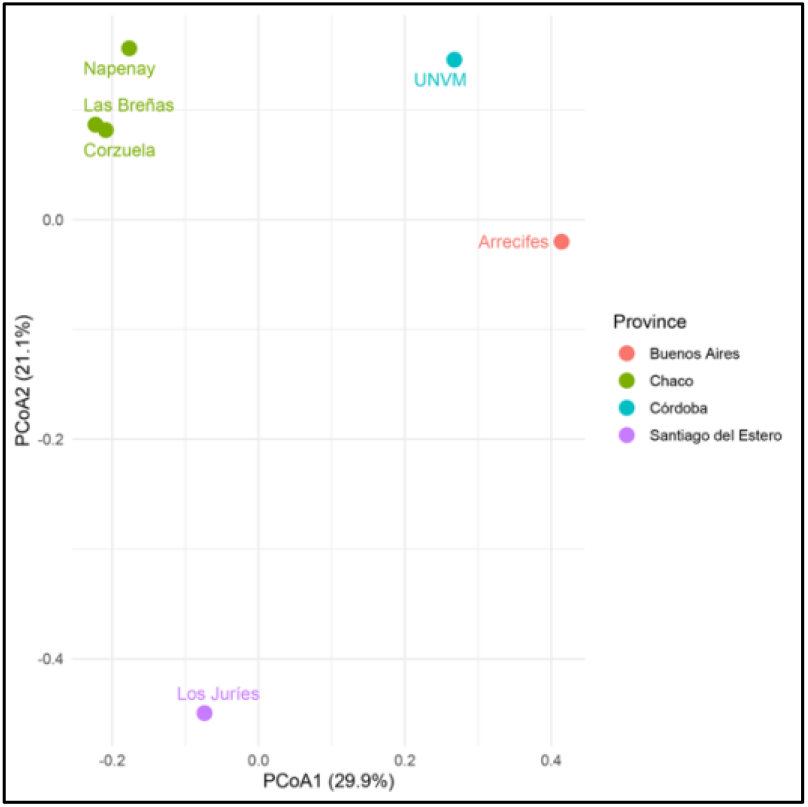
Beta diversity of soil bacterial communities based on Bray–Curtis dissimilarity. Principal coordinate analysis (PCoA) showing clustering of samples according to geographic origin. Samples from the Chaco region grouped together, whereas samples from other regions showed clear separation.

**Figure 5.**
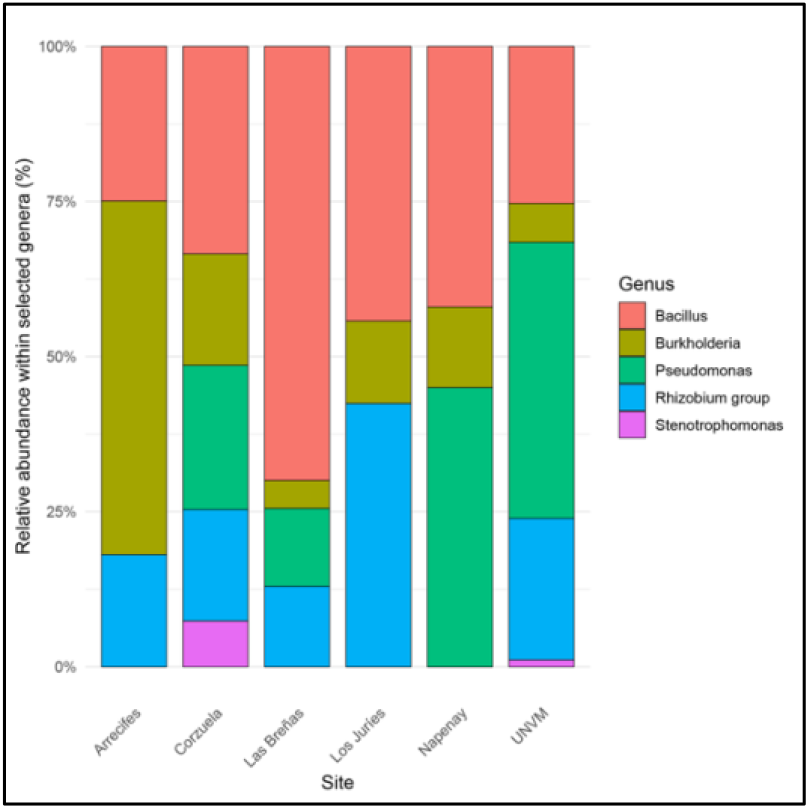
Distribution of agronomically relevant bacterial genera across soil samples. Relative abundance of selected plant-beneficial and plant-associated genera across sampling sites. *Bacillu*s and *Pseudomonas* were among the most abundant genera, with variation across sites. Genera were selected based on their reported roles in plant growth promotion and biological control.

Samples from the Chaco region (Corzuela, Las Breñas, and Napenay) clustered closely together, indicating a similar microbial composition. In contrast, samples from Arrecifes (Buenos Aires), Los Juríes (Santiago del Estero), and UNVM (Córdoba) were clearly separated from the Chaco cluster along the first two axes. The first and second axes explained 29.9% and 21.1% of the total variance, respectively, highlighting substantial differences in community structure across sites. Overall, these results suggest that geographic location and local environmental conditions are key factors shaping soil microbial community composition.

### Distribution of agronomically relevant genera

The most abundant genera identified across samples included *Bradyrhizobium, Bacillus, Rhizobium* and *Pseudomonas*, all of which are known to play important roles in agricultural systems. *Bradyrhizobium* and *Rhizobium* (rhizobia) were particularly abundant in Corzuela, Napenay, Los Juríes and Arrecifes. These genera are well known for their ability to establish symbiotic relationships with legumes, contributing to biological nitrogen fixation and improving soil fertility.

*Bacillus* was dominant in several samples, including Las Breñas and UNVM. This genus includes species such as *Bacillus subtilis* and *Bacillus thuringiensis*, which are widely used in agriculture as plant growth-promoting rhizobacteria (PGPR) and biological control agents against plant pathogens and insect pests.

*Pseudomonas* was detected in all samples, with higher relative abundance in UNVM. This genus includes both plant-beneficial species and plant-associated pathogens, highlighting its dual ecological role. Minor genera with agronomic relevance, including *Burkholderia, Stenotrophomonas, Herbaspirillum, Ralstonia* and *Xanthomonas*, were also detected at lower abundance. Notably, several important plant pathogenic genera, including *Enterobacter, Agrobacterium, Erwinia, Xylella, Dickeya* and *Pectobacterium*, were not detected in any of the analyzed samples.

### Functional groups and biocontrol-related taxa

Genera associated with potential biocontrol and potential entomopathogenic activity were detected across all samples, including *Bacillus* and *Pseudomonas*, with additional genera such as *Paenibacillus, Lysinibacillus* and *Brevibacillus* detected at lower abundance.

*Bacillus* was among the most prevalent genera across sites, whereas UNVM showed a higher relative representation of *Pseudomonas* and *Paenibacillus*, indicating spatial variability in the distribution of taxa with potential biocontrol activity.

Overall, these results indicate that the analyzed soils harbor diverse microbial taxa with potential relevance for sustainable agriculture and biotechnological applications, although functional roles cannot be reliable inferred from 16S rRNA data.

## Discussion

The present study provides an exploratory characterization of soil bacterial communities from agroecosystems located in north-central Argentina using 16S rRNA amplicon sequencing. Overall, the results revealed highly diverse and structured microbial communities, with clear differences associated with geographic location and local environmental conditions.

The dominance of Proteobacteria and Actinobacteriota across all samples is consistent with previous studies in agricultural soils, where these phyla are typically associated with copiotrophic lifestyles and nutrient-rich environments (Dai et al., 2018; Fierer et al., 2012). Their prevalence has been linked to fertilization practices, particularly nitrogen inputs, which promote fast-growing microbial groups (Dai et al., 2018). In contrast, the presence of Acidobacteriota, especially in Las Breñas, suggests the coexistence of oligotrophic taxa adapted to lower nutrient availability, as previously described for this phylum (Kielak et al., 2016).

Alpha diversity analysis indicated consistently high bacterial diversity across all samples, with Shannon index values comparable to those reported in other agricultural soils (Carbonetto et al., 2014; Fierer et al., 2013). Although no statistical comparisons were performed due to the limited number of samples, trends suggest that sites such as UNVM and Arrecifes harbor more complex and evenly distributed microbial communities.

Beta diversity analysis further supported the influence of geographic and environmental factors on microbial community structure. The clustering of samples from Chaco indicates similar microbial composition within this region, whereas samples from other regions displayed distinct community profiles. These findings are consistent with previous studies showing that soil properties, climate, and land use are major drivers of microbial community assembly in agricultural systems (Kraut-Cohen et al., 2020).

At the genus level, several taxa with well-established agronomic relevance were identified, including *Bradyrhizobium, Rhizobium, Bacillus* and *Pseudomonas*. The high abundance of rhizobia-related genera is consistent with the cultivation of legumes such as soybean and reflects the importance of biological nitrogen fixation in these systems (Amaresan et al., 2020). Likewise, the widespread presence of *Bacillus* supports its recognized role as a plant growth-promoting rhizobacterium (PGPR) and biological control agent (Backer et al., 2018; Zhang et al., 2020).

The genus *Pseudomonas* was detected across all samples, reinforcing its ecological versatility. This genus includes both plant-beneficial and plant-associated pathogenic species, highlighting the functional complexity of soil microbial communities (Mansfield et al., 2012; Qessaoui et al., 2019).

Additionally, several genera associated with potential biocontrol and entomopathogenic activity were identified, including *Bacillus, Paenibacillus, Lysinibacillus, Brevibacillus* and *Pseudomonas*. Members of these genera are widely recognized for their roles in biological control through mechanisms such as toxin production, antibiosis, and induction of plant systemic resistance (Bais et al., 2004; Palma et al., 2014; Ramos et al., 2014). The differential abundance of these taxa among sites suggests variability in the potential for natural biological control across agroecosystems. However, it is important to note that 16S rRNA amplicon sequencing does not provide direct functional information or species-level resolution, limiting the ability to accurately infer ecological roles (Dubey et al., 2019).

Despite these insights, nitrifying bacteria were not prominently detected at the genus level, likely due to their low abundance or limitations in taxonomic resolution inherent to 16S rRNA sequencing (Dubey et al., 2019; Prosser et al., 2007). Future studies using targeted approaches or shotgun metagenomics may provide a more accurate assessment of functional groups involved in nitrogen cycling.

Taken together, the observed patterns are consistent with previous studies conducted in managed agricultural systems, where microbial community composition is shaped by a combination of environmental variables, soil properties, and agricultural practices (Fierer et al., 2012; Kraut-Cohen et al., 2020). The results presented here contribute to expanding current knowledge of soil microbiomes in Argentine agroecosystems beyond the Pampas region and provide a baseline for future large-scale metagenomic studies.

## Conclusions

The results obtained in this study contribute to expanding current knowledge of soil bacterial communities in agricultural systems from Argentine regions outside the Pampas, which remain relatively underexplored. Although the data presented here are exploratory, they provide a baseline for future high-throughput metagenomic studies involving a larger number of samples across diverse agroecosystems in Argentina. Collectively, the observed microbial community patterns are consistent with those reported in other managed agricultural systems, supporting the role of environmental and agricultural management factors in shaping soil microbiomes.

## Author contributions

Conceptualization, L.P.; methodology, A.L.G., A.M., C.P., E.E.D.V., L.C. and L.P.; formal analysis, A.L.G., A.M., C.P., E.E.D.V., L.C. and L.P.; investigation, A.L.G., A.M., C.P., E.E.D.V., L.C. and L.P.;resources, L.P.; data curation, L.P.; writing—original draft preparation, L.P.; writing—review and editing, L.P.; visualization, L.P.; supervision, L.P.; project administration, L.P.; funding acquisition, L.P. All authors have read and agreed to the published version of the manuscript.

## Data availability

The raw 16S rRNA gene sequencing data generated in this study are publicly available in the NCBI Sequence Read Archive (SRA) under BioProject accession PRJNA722672.

## Acknowledgements

Leopoldo Palma gratefully acknowledges the Spanish Ministry of Science, Innovation, and Universities, the Spanish State Research Agency, and the European Union for funding his Ramón y Cajal contract (grant ref. RYC2023-043507-I).

## References

Aizen, M. A., et al., 2009. Expansión de la soja y diversidad de la agricultura argentina. Ecología austral. 19, 45–54.

Ali, B., 2019. Functional and genetic diversity of bacteria associated with the surfaces of agronomic plants. Plants. 8, 91.

Amaresan, N., et al., 2020. Beneficial microbes in agro-ecology: bacteria and fungi. Academic Press.

Backer, R., et al., 2018. Plant growth-promoting rhizobacteria: context, mechanisms of action, and roadmap to commercialization of biostimulants for sustainable agriculture. Frontiers in plant science. 9, 1473.

Bais, H. P., et al., 2004. Biocontrol of Bacillus subtilis against infection of Arabidopsis roots by Pseudomonas syringae is facilitated by biofilm formation and surfactin production. Plant physiology. 134, 307–319.

Bran, D., et al., 2017. Los indicadores de calidad de suelo como un componente de la sustentabilidad de los agroecosistemas. WILSON, Marcelo Germán. Manual de indicadores de calidad del suelo para las ecorregiones de Argentina. Entre Ríos. INTA. 15–17.

Carbonetto, B., et al., 2014. Structure, composition and metagenomic profile of soil microbiomes associated to agricultural land use and tillage systems in Argentine Pampas. PLoS One. 9, e99949.

Dai, Z., et al., 2018. Long-term nitrogen fertilization decreases bacterial diversity and favors the growth of Actinobacteria and Proteobacteria in agro-ecosystems across the globe. Global change biology. 24, 3452–3461.

Dubey, A., et al., 2019. Soil microbiome: a key player for conservation of soil health under changing climate. Biodiversity and Conservation. 28, 2405–2429.

Fierer, N., et al., 2013. Reconstructing the microbial diversity and function of pre-agricultural tallgrass prairie soils in the United States. Science. 342, 621–4.

Fierer, N., et al., 2012. Comparative metagenomic, phylogenetic and physiological analyses of soil microbial communities across nitrogen gradients. Isme j. 6, 1007–17.

Geisseler, D., Scow, K. M., 2014. Long-term effects of mineral fertilizers on soil microorganisms–A review. Soil Biology and Biochemistry. 75, 54–63.

Guimarães, R. P., 1998. Aterrizando un cometa: indicadores territoriales de sustentabilidad.

Kielak, A. M., et al., 2016. The Ecology of Acidobacteria: Moving beyond Genes and Genomes. Front Microbiol. 7, 744.

Kraut-Cohen, J., et al., 2020. Effects of tillage practices on soil microbiome and agricultural parameters. Science of the Total Environment. 705, 135791.

Leinonen, R., et al., 2010. The sequence read archive. Nucleic acids research. 39, D19–D21.

Mansfield, J., et al., 2012. Top 10 plant pathogenic bacteria in molecular plant pathology. Molecular plant pathology. 13, 614–629.

Morello, J., et al., 2012. Ecorregiones y complejos Ecosistémicos de Argentina. Orientación Gráfica Editora, Buenos Aires.

Oksanen, J., et al., 2013. Community ecology package. R package version. 2, 321–326.

Palma, L., et al., 2014. Bacillus thuringiensis toxins: an overview of their biocidal activity. Toxins (Basel). 6, 3296–3325.

Prosser, J. I., et al., 2007. The role of ecological theory in microbial ecology. Nature Reviews Microbiology. 5, 384–392.

Qessaoui, R., et al., 2019. Applications of new rhizobacteria Pseudomonas isolates in agroecology via fundamental processes complementing plant growth. Scientific reports. 9, 12832.

Ramos, J.-L., et al., 2014. Pseudomonas: Volume 7: New aspects of Pseudomonas biology. Springer.

Rascovan, N., et al., 2013. The PAMPA datasets: a metagenomic survey of microbial communities in Argentinean pampean soils. Microbiome. 1, 21.

Zhang, X., et al., 2020. Applications of Bacillus subtilis spores in biotechnology and advanced materials. Applied and environmental microbiology. 86, e01096–20.

